# Behavioral responses of free flying *Drosophila melanogaster* to shiny, reflecting surfaces

**DOI:** 10.1101/2022.12.30.522304

**Authors:** Thomas Mathejczyk, Edouard J. Babo, Erik Schönlein, Nikolai V. Grinda, Andreas Greiner, Nina Okrožnik, Gregor Belušič, Mathias F. Wernet

## Abstract

Active locomotion plays an important role in the life of many animals since it permits to explore the environment and find vital resources. Most insect species rely on a combination of visual cues such as celestial bodies, landmarks, or linearly polarized light to navigate or to orient themselves in their surroundings. In nature, linearly polarized light can arise either from atmospheric scattering or from reflections off shiny non-metallic surfaces like water or shiny foil. Although multiple reports described different behavioral responses of various insects to such shiny surfaces, little is known about the retinal detectors or the underlying neural circuits. Our goal was to quantify the behavioral responses of free flying *Drosophila melanogaster*, a molecular genetic model organism that allows for systematic dissection of neural circuitry. Fruit flies were placed in a custom-built arena with controlled environmental parameters (temperature, humidity, and light intensity). Flight densities and landings were quantified for hydrated and dehydrated fly populations when separately exposed to three different stimuli such as a diffusely-reflecting matt plate, a small patch of shiny foil, versus real water. Our analysis reveals for the first time that flying fruit flies indeed use vision to guide their flight maneuvers around shiny surfaces.

## Introduction

The ability to navigate is one of the most prominent locomotive achievements of insects (Heinze 2017). This requires animals to perceive their position within their habitat (allowing them to orient) and/or to integrate this signal with spatial memory in the case of returning to their previous location (leading to navigation) (Collett, Chittka et al. 2013, Heinze 2017). To navigate or orient, insects combine and integrate diverse sensory cues which provide them with information about their surroundings, and consequently induce an adaptive behavior such as following a trajectory (Heinze, 2017). Many species have evolved integrated navigational mechanisms, relying on global signals in the sky and/or local signals like landmarks (Collett, Lent et al. 2014, El Jundi, Foster et al. 2016), wind direction (Dacke, Bell et al. 2019), and idiothetic signals like optic flow (Mauss and Borst 2020) and proprioception (Wittlinger, Wehner et al. 2006).

The celestial pattern of linearly polarized skylight is used by many insect species for improving their orientation/navigation skills (Heinze, 2017)(Wehner and Labhart 2006). The e-vectors of skylight scattered in the atmosphere create a polarization pattern across the sky, which changes according to the location of the sun (Wehner 2001, Heinze 2017, Mathejczyk and Wernet 2017). Many insects have evolved specialized visual systems, including specialized retinal detectors as well as underlying circuitry, to use this pattern for navigation and orientation (Labhart and Meyer 1999, Homberg 2015, Dacke and El Jundi 2018, Sancer, Kind et al. 2019, Kind, Belušič et al. 2020, Sancer, Kind et al. 2020). Importantly, sunlight also becomes linearly polarized through reflections off shiny surfaces like water, animal or plant cuticles, or any nonmetallic shiny surfaces (Wehner, 2001). Most of these manifest angles of polarizations parallel to the reflecting surface and reach their maximum at Brewster’s angle (53° from normal for air-water interface) (Wehner 2001). Although many behavioral responses have been reported, much less is known about how insects detect polarized reflections (Mathejczyk and Wernet 2017, Heinloth, Uhlhorn et al. 2018, Yadav and Shein-Idelson 2021) and the retinal detectors of horizontally polarized reflections remain largely unclear, with only very few exceptions (Schneider and Langer 1969, Schwind 1983, Meglic, Ilic et al. 2019).

The *Drosophila melanogaster* retina is composed of approximately 800 ommatidia each containing eight light-sensitive sensory cells or photoreceptors (named R1-R8). Six outer photoreceptors surround the inner photoreceptors R7 and R8 (Kind, Longden et al. 2021). Based on the molecular and physiological characteristics of inner photoreceptors, ommatidia can be divided into at least three groups, called pale, yellow, and DRA (Wernet, Perry et al. 2015, Kind, Longden et al. 2021). Together, pale and yellow ommatidia form the genetically specified color vision detection system of the fly retina and are randomly distributed at an uneven ratio with 30% pale and 70% yellow (Feiler, Bjornson et al. 1992, Wernet, Mazzoni et al. 2006, Bell, Earl et al. 2007). Only at the dorsal-most edge of the fly retina, in the so-called dorsal rim area (DRA), inner photoreceptors R7 and R8 from the same ommatidia form homochromatic polarization sensors with orthogonal e-vector tunings of R7 vs R8 (Wada 1974, Fortini and Rubin 1990, Wernet, Labhart et al. 2003). For *Drosophila* it was shown that DRA ommatidia are both necessary and sufficient to detect linearly polarized skylight (Wernet, Velez et al. 2012). Surprisingly, *Drosophila* can also perceive linearly polarized light with the ventral half of its retina (Wolf, Gebhardt et al. 1980, Wernet, Velez et al. 2012, Velez, Gohl et al. 2014). Thus, linearly polarized light is not only important for navigation but might also serve the detection of other visual signals on the ground.

Reflected polarized light can provide crucial and reliable detail for the detection of a variety of salient stimuli, like water bodies, shiny food, insect cuticles, or even an appropriate place for oviposition (Mathejczyk and Wernet 2017, Heinloth, Uhlhorn et al. 2018, Yadav and Shein-Idelson 2021). Such ventral detection of linearly polarized light has been shown in a variety of species, ranging from locusts (Shashar, Sabbah et al. 2005) and hemipterans (Schwind 1984), to dragonflies (Wildermuth 1998), chironomids (Lerner, Meltser et al. 2008), and mayflies (Farkas, Szaz et al. 2016). Furthermore, it has been shown that shiny non-metallic surfaces evoke behavioral responses in semi-aquatic insects, equivalent to what water does (Kriska, Bernath et al. 2007, Farkas, Szaz et al. 2016). The behavioral response to reflected polarized light can be attractive, like in the case of the backswimmer *Notonecta glauca*, which manifests a diving reflex when presented to a polarized surface (Schwind 1984, Schwind 1985), or it can be repulsive, like in case of the desert locust *Schistocerca gregaria*, which avoid linearly polarized surfaces, presumably in order to avoid drowning (Shashar, Sabbah et al. 2005). Even though all these species appear to share the ability to perceive polarized reflections emanating from the ventral side, there exists no obvious common anatomical substrate now obvious mechanisms for this purpose among the insects (Heinloth et al., 2018).

Here we investigated ventral perception of linearly polarized light by unrestrained, flying *Drosophila melanogaster*. Although well studied in tethered flight for both dorsally and, in some cases, ventrally presented stimuli (Wolf, Gebhardt et al. 1980, Weir and Dickinson 2012, Velez, Gohl et al. 2014, Velez, Wernet et al. 2014, Warren, Weir et al. 2018, Mathejczyk and Wernet 2019, Mathejczyk and Wernet 2020), no experiments addressing the behavioral responses of free-flying *Drosophila* being ventrally exposed to reflected linear polarized light have yet been conducted. Furthermore, *Drosophila* can be both repelled by, as well as attracted to water, depending on their physiological state (hydrated vs dehydrated) (Ji and Zhu 2015). Hence, we asked whether free-flying *Drosophila* would manifest a specific behavioral response when ventrally exposed to linearly polarized reflections.

## Materials and Methods

### Flies

All experiments were conducted using four-day-old male and female *Drosophila melanogaster* (strain Top Banana) provided by Eugenia Chiappe (personal donation). Flies were raised on standard medium at constant temperature (25°C) and humidity (50%-60%) within a 12/12 light/dark cycle. For desiccation, flies were transferred into vials containing desiccant (Drierite®) at the bottom (about 1cm fill height) covered by a small ball of cotton wool to avoid any contact of the flies with the desiccant. Flies were desiccated for six hours with the desiccation vials placed in the incubator (25°C, 50-60% humidity). Due to logistical reasons, all experiments involving dehydrated flies were at their night activity peak (approximately 2 hours before their subjective “sunset”).

### Experimental setup

To test behavioral responses towards shiny/linearly polarizing surfaces during flight, we designed and constructed a custom behavioral assay. In this assay, flies were transferred into a white plastic bucket (diameter approximately 30cm) with the bottom plate cut off and filmed (Point Grey® Blackfly BFLY-PGE-2356M + Lee Filters 87C infrared pass filter) from above against near-infrared illumination (LED array, 850nm) coming from below. White cloth was attached to the top of the arena to prevent flies from escaping during experiments. Visible (white light) illumination was provided using RGB LED strips (12V at 0.1A total) at the top of the arena for which the red, green and blue channels were calibrated iso-quantally using a photo-spectrometer (Ocean Optics Flame UV-VIS). A matt (surface manually roughed up using sandpaper) white acrylic plate (320 mm x 320 mm, 3 mm thick) served as a weakly polarizing arena bottom for visual light coming from above, as well as a diffuser for the near-infrared illumination coming from below. The assay was placed within a black metal enclosure for optical shielding and climate control. The temperature was up-regulated using a carbon heater (#100575, termowelt.de) placed at the bottom of the metal enclosure and an external PID-controller (RT4-121-Tr21Sd, pohltechnic.com). We constructed 2 small humidifiers by placing small fans (40 mm x 40 mm) on top of cut plastic bottles with ventilation holes (filled with water + pieces of replacement humidifier filter Philips 2000 HU4801/02). The humidifiers were controlled using an external hygrostat.

### Visual stimuli

We tested three different visual stimuli presented at the bottom of the arena: an all-matt acrylic plate showing an overall low degree of linear polarization (Dolp) of reflected light, an all-matt acrylic plate with a shiny, highly reflective center (60mm diameter clear overhead projection foil, manufacturer and brand: 1 CLASSIC 6070 overhead transparencies) and an all-matt acrylic plate with a 60mm diameter, 1mm deep (sloped walls) indentation in the center, filled with water.

### Polarimetry

Polarimetric and intensity measurements were acquired using a polarimetry camera (PHX050S1-PC, Lucid Vision Labs). To image the center of the arena from different horizontal angles, a tripod in combination with an electronic level was used. For these measurements, we also replaced the white bucket with cut-in-half bucket of the same type. This allowed for imaging the arena from a variety of angles, as well as capturing reflections off of the bucket walls, as would also be present during experiments. Polarimetric and intensity images were saved as 8-bit greyscale TIFF files and the mean Dolp and intensity for zone 1 and 2 were quantified using FIJI (https://imagej.net/software/fiji/).

### Experimental procedure and data acquisition

All experiments were conducted at 30°C and 50-60% relative humidity. Before each experiment, approximately 100 flies were transferred into empty vials and counted manually. Flies were introduced to the arena by inserting the vial through a slit between the white cloth and the bucket and letting flies enter on their own. Each experiment was recorded for 60 minutes (43fps, 8bit greyscale, 580×580pixel resolution) and videos were saved using MJpeg compression. Each experimental condition was at least recorded eight times and after each recording, the flies were vacuumed out of the arena.

### Tracking of flight behavior

In order to separate flying from walking flies and to track their positional data over time, we developed a custom Matlab script. From the recorded videos the code computes intensity differences between consecutive frames. At the framerate at which the videos were recorded (43fps), intensity differences between consecutive frames for walking flies are very low due to their relatively slow walking speed. However, due to the higher flight velocities, flying flies usually move over a larger distance than their body length between consecutive frames, resulting in large intensity differences at the position where a flying fly ‘appears’ in a consecutive frame. These intensity differences were filtered, thresholded and ultimately used to locate the X and Y coordinates of flight occurrences for many flies simultaneously. This method allows for quick quantification of probabilistic distributions of flight occurrences in large fly populations without having to keep the tracking identities of individual flies for discriminating between walking and flight. To verify the robustness of this method, 8 hours of videos containing the thresholded files were compared frame by frame to the original recordings. Depending on the flight angle, in some cases (e.g. a fly flying directly towards the camera) flight activity could not be detected as such due to low intensity differences between consecutive frames, resulting in a rather conservative estimate of flight activity. However, due to the large number of flies and observations, overall flight activity is represented very robustly using this method.

### Tracking of landings

Landings were tracked manually using FIJI (https://imagej.net/software/fiji/). For tracking, every ten consecutive frames of the original video were converted into a minimal intensity projection, allowing for better visibility of flight trajectories and also an increased tracking speed. Whenever a landing was observed, a point marker was placed, saving the X and Y coordinates of each landing. Due to the very high amount of landings of thirsty flies, for matt thirsty and shiny foil thirsty recordings we only tracked the first 1000 landings per video.

### Statistical Analysis

Statistical analysis was performed using Matlab. To calculate the distribution probability for flight and landing detections, we divided the arena into 5 concentric zones, with the center zone having a diameter of 60mm (same as shiny foil and water stimulus) and each consecutive zone having a 60mm larger diameter than the next smaller area. Next, the number of flight detections and landings was calculated for each zone and normalized for the area of the respective zones in order to finally calculate the normalized detection probability in %. Normalized detection probabilities between zone 1 and zone 2 were compared using the two-sample t-test for flights and landings, respectively. For flight and landing statistics, we counted one hour of video recording as one sample (n=1). The number of tested flies and detections for each recorded video and condition is described in detail in supplementary table 1.

## Results

The aim of this study was to determine whether freely flying flies would interact with shiny and therefore potentially linearly polarizing surfaces, by analyzing both flight trajectories as well as the distribution of landings.

### Behavioral setup

We built a novel assay allowing for probabilistic quantifications of spatial flight distributions (Fig. 1). Freely moving populations of wild type flies were filmed from above against a near-infrared illumination within a temperature- and humidity-controlled cylindrical arena (Fig. 1a). To test the flies’ behavioral responses towards either one of three different stimuli, the bottom of the arena was either completely matt (rough surface), or it was equipped either with an additional small piece of shiny overhead projector foil in the center, or with a small body of water at the same location (Fig. 1b). We also developed a computationally fast and easy-to-use procedure of automatically distinguishing flying from walking flies based on intensity differences of consecutive video frames, allowing for the quantification of probabilistic spatial distributions during flight (Fig. 1c) (see materials and methods).

**Figure 1:**
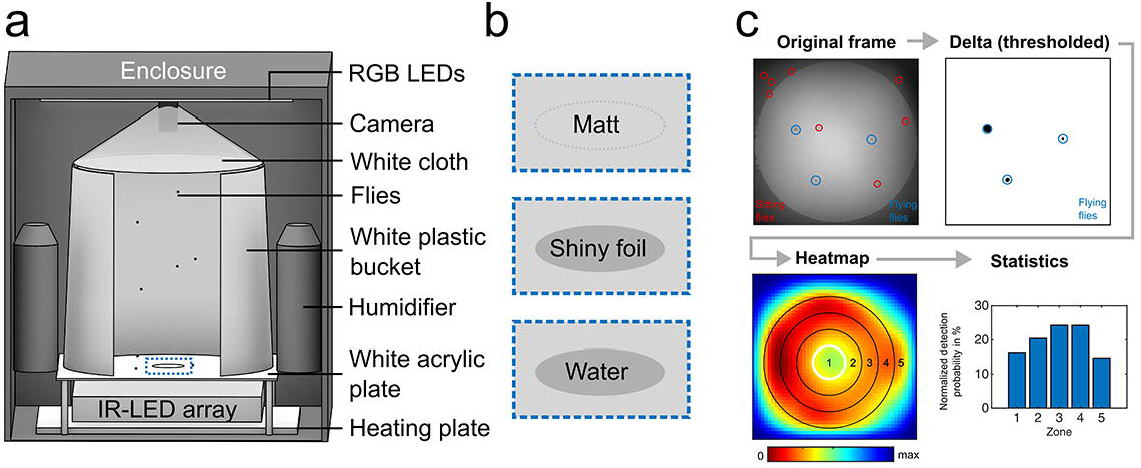
Data acquisition and processing. (a) Schematic of the flight arena setup with all main components (see material & methods). (b) Summary of the three stimuli used as bottom plates (see material & methods). (c) Flow chart of the flight density tracking procedure (see material & methods).

### Stimulus characterization

Polarimetric characterization of the three different visual stimulus conditions used here was performed as previously described (Meglic, Ilic et al. 2019), and revealed relatively low values for the degree of linear polarization (Dolp) of around 15%, across all tested elevation angles for both the matt and water stimulus condition and also for the matt region surrounding the shiny overhead foil (zone 2), across all stimulus conditions (Fig.2 a, b, c). As expected, an elevated degree of linear polarization of around 25% was measured only when the shiny overhead foil was placed in the center of the matt plate (Fig. 2b). Simultaneously performed intensity measurements revealed only little intensity differences between the centrally located zone 1 and the surrounding zone 2 for the matt and water stimulus condition whereas the intensity of the shiny foil center was even lower than its’ surrounding zone 2 at low elevation angles and slightly higher only at 60° elevation (Fig.2 d-e). Hence, a shiny overhead foil placed at the center of the arena was from most fly aspects not significantly brighter than the surrounding matt surface.

**Figure 2:**
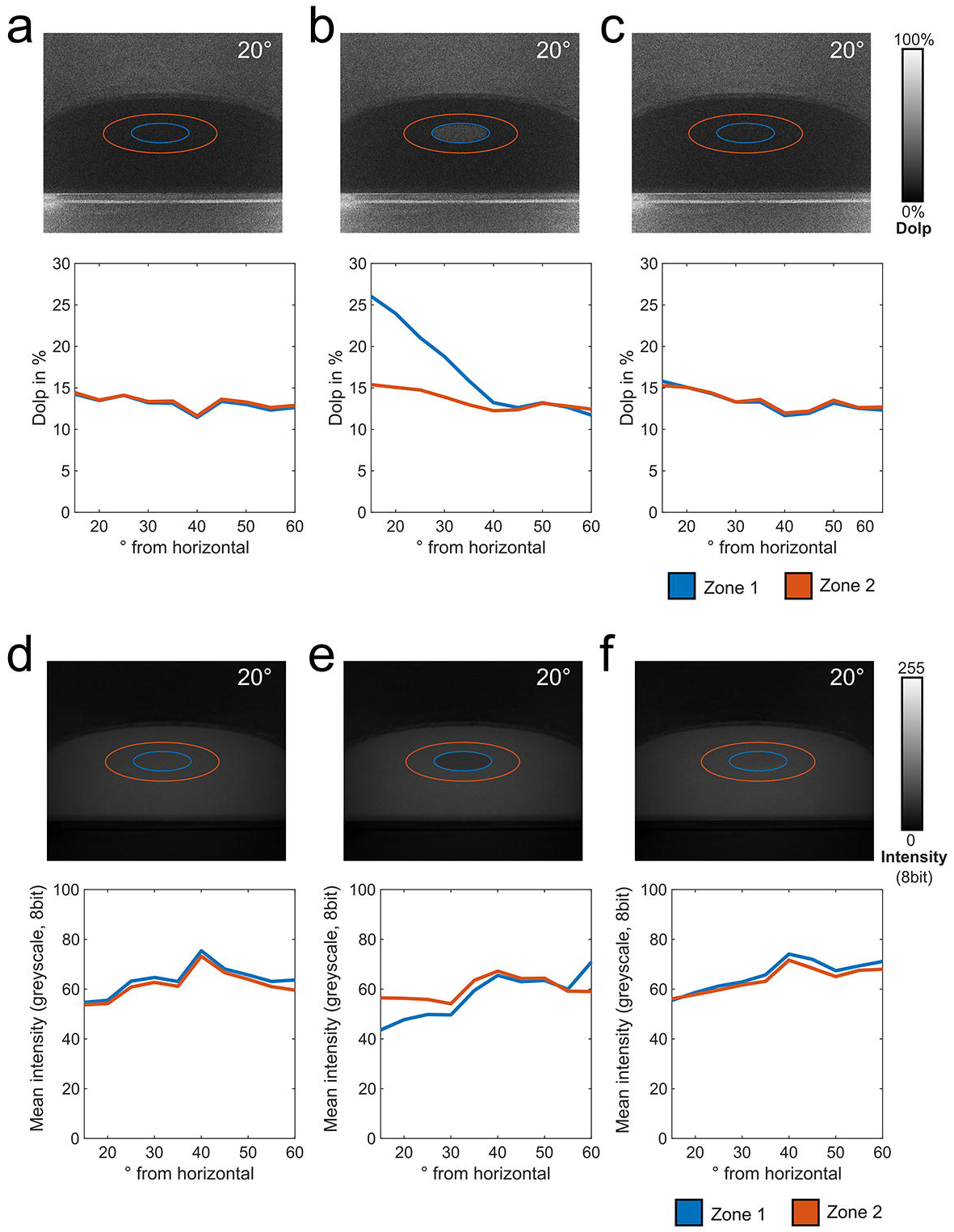
Polarimetric characterization of the stimuli used. (a)-(c) Polarimetric quantification of the degree of linear polarization (Dolp) of the center (zone 1) versus immediate surround (zone 2) for the three stimulus conditions used across different incident angles (see material & methods). (e)-(f) Quantification of main intensity of the center (zone 1) versus immediate surround (zone 2) for the three stimulus conditions used across different incident angles.

### Flight behavior

For both matt and water stimuli, we found only weak differences in flight detection probability between zone 1 (center) and zone 2 (surround) in thirsty as well as in not-thirsty fly populations (Fig.3 a, c, d, f). In contrast, the same analysis revealed a strongly reduced flight detection probability above zone 1 when the shiny overhead foil was placed there, as compared to its surrounding matt zone 2 (Fig. 3b). This effect was even more pronounced in thirsty flies (Fig. 3e). Differences in flight detection probability between zone 1 and zone 2 were significantly larger with the shiny foil in the center when directly compared to the matt and water stimulus, respectively (Fig.4a) but did not differ significantly between thirsty and non-thirsty flies.

**Figure 3:**
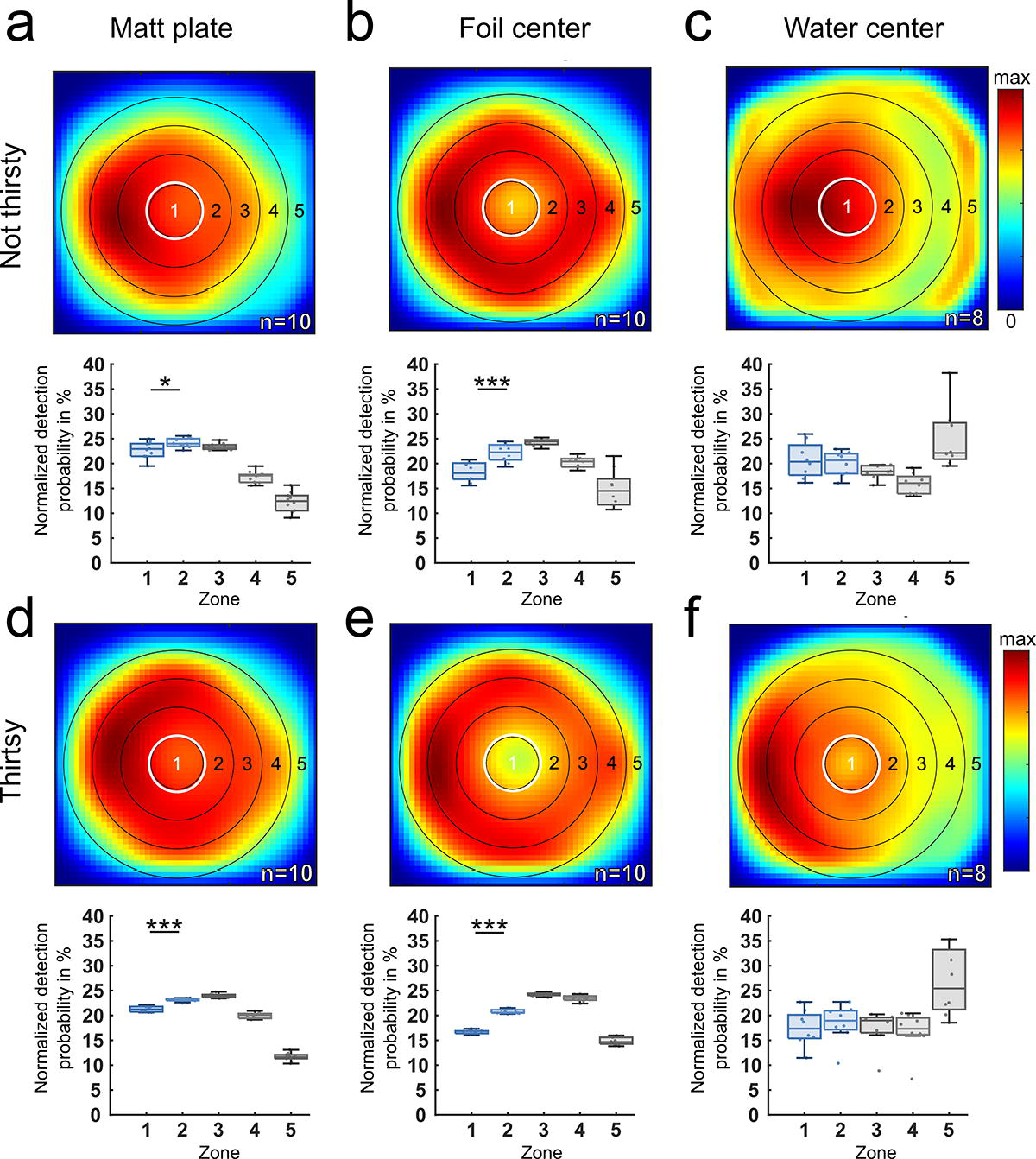
Flight densities of hydrated vs dehydrated flies in response to different stimuli. Summary and direct comparison of flight densities of hydrated flies (a-c) versus dehydrated flies (d-f) in the arena after 60 minutes of recording of three experimental conditions: (a), (d) completely matt surface; (b), (e) matt plate with a shiny foil center; (c), (f) matt plate with a water center. A heatmap of false-colored flight densities is shown, as well as the normalized detection probability for flying *Drosophila* to be in each concentric zone. n indicates the number of 60-minute recordings. *p<0.05, **p<0.01, ***p<0.001.

**Figure 4:**
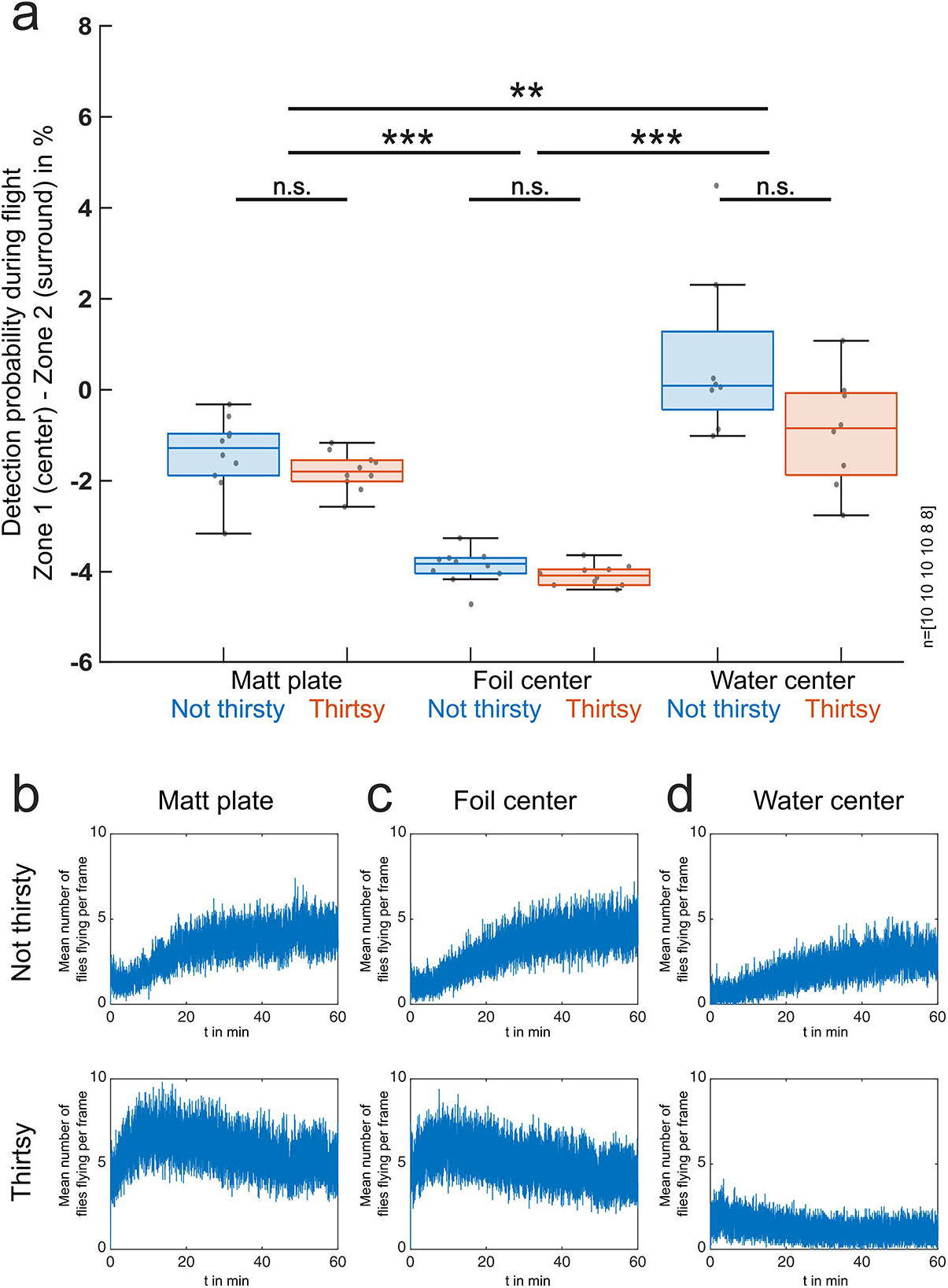
Statistics of hydrated vs dehydrated flies and overall flight activity. (a) Statistical summary of detection probability during flight (zone 1 – zone 2), directly comparing hydrated and dehydrated flies in all three experimental conditions. Number of 60-minute recordings from left to right indicated at the right. *p<0.05, **p<0.01, ***p<0.001. (b) – (d) Summary and direct comparison of flight activity (mean number of flies flying per frame) for hydrated flies (top) versus dehydrated flies (bottom) in the arena after 60 minutes of recording of three experimental conditions: (b) completely matt surface; (c) matt plate with a shiny foil center; (d) matt plate with a water center.

Calculating the average number of flying flies per frame over time allowed us to also quantify flight activity dynamics for both thirsty and not-thirsty flies, under all stimulus conditions. When presented with a matt or shiny center, respectively, average flight activity increased over time in non-thirsty flies, whereas thirsty flies showed high flight activity from the beginning of the experiment (Fig. 4 b-d). However, when presented with water in the center, both thirsty and non-thirsty flies, always showed relatively low flight activity. Visual inspection revealed that many of them were often sitting right adjacent to the water surface, over long periods of time (see supplemental Fig. S1).

### Distribution of fly landings

Similar to our flight probability observations, the manual quantification of fly landings revealed only weak differences between landing probability in zone 1 (center) versus zone 2 (surrounding) for the overall matt stimulus condition, both in thirsty as well as in not-thirsty flies (Fig.5a,d). In contrast, we observed a strongly reduced landing probability in zone 1 both when the shiny overhead foil was placed there, as well as in the case of a real water body. This effect persisted for both thirsty and not-thirsty flies (Fig.5b, c, e, f). A direct comparison across experiments revealed that the differences in landing probability between zone 1 and zone 2 were significantly larger in those experiments using shiny overhead foil or water stimuli in zone 1, as compared to the experiments where both zones 1 and 2 were matt (Fig. 6). Differences in landing probabilities for zone 1 and zone 2 were significant between thirsty and non-thirsty flies only when shiny foil was used as a stimulus in the center.

**Figure 5:**
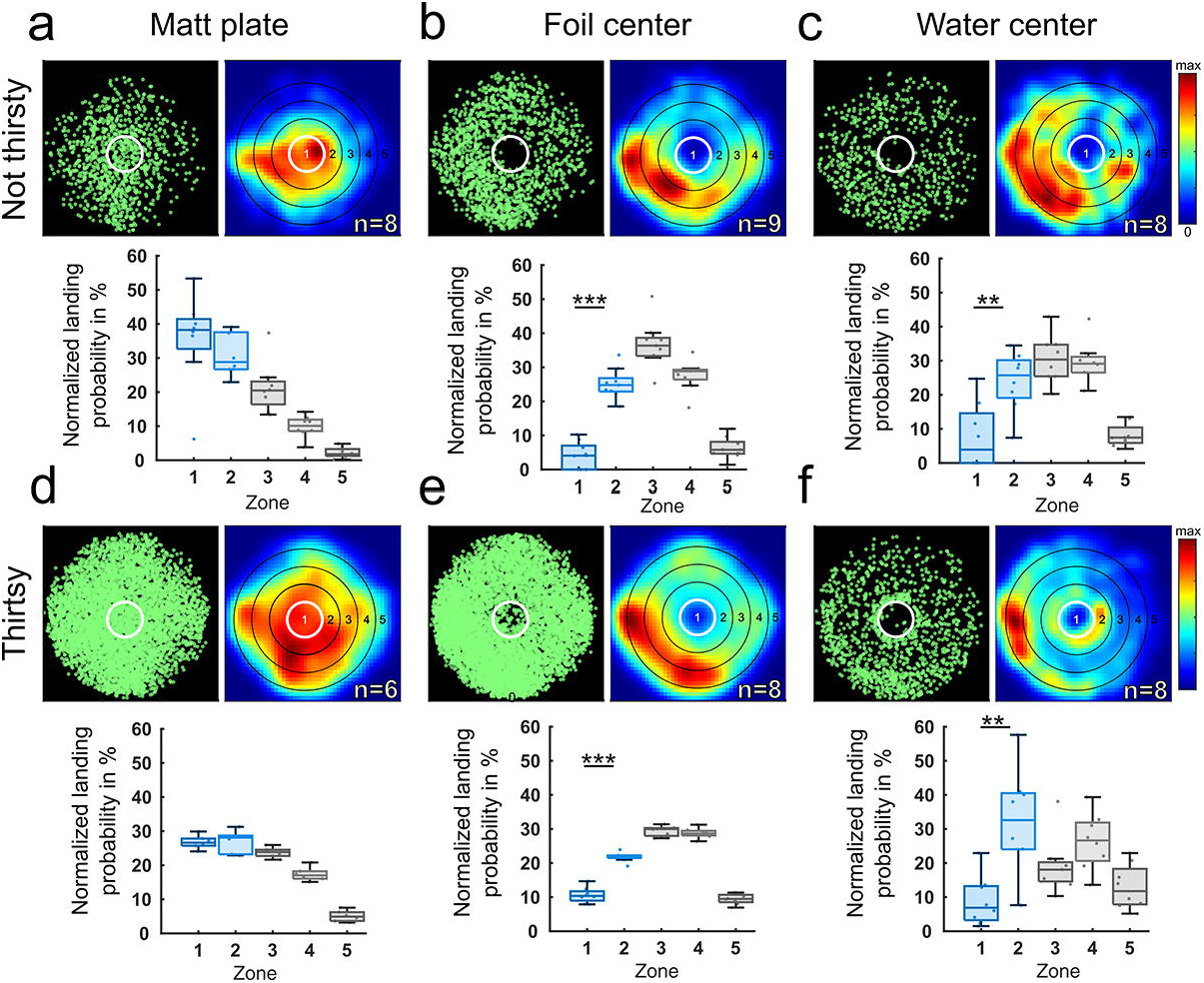
Quantification of landings of hydrated vs dehydrated flies in response to three stimuli. Summary and direct comparison of landing events of hydrated flies (a-c) versus dehydrated flies (d-f) in the arena after 30 minutes of recording of three experimental conditions: (a), (d) completely matt surface; (b), (e) matt plate with a shiny foil center; (c), (f) matt plate with a water center. For each condition, manually tracked landings are shown (green), as well as a heatmap of false-colored landing densities, and the normalized detection probability for landing *Drosophila* to be in each concentric zone. n indicates the number of 60-minute recordings. *p<0.05, **p<0.01, ***p<0.001.

**Figure 6:**
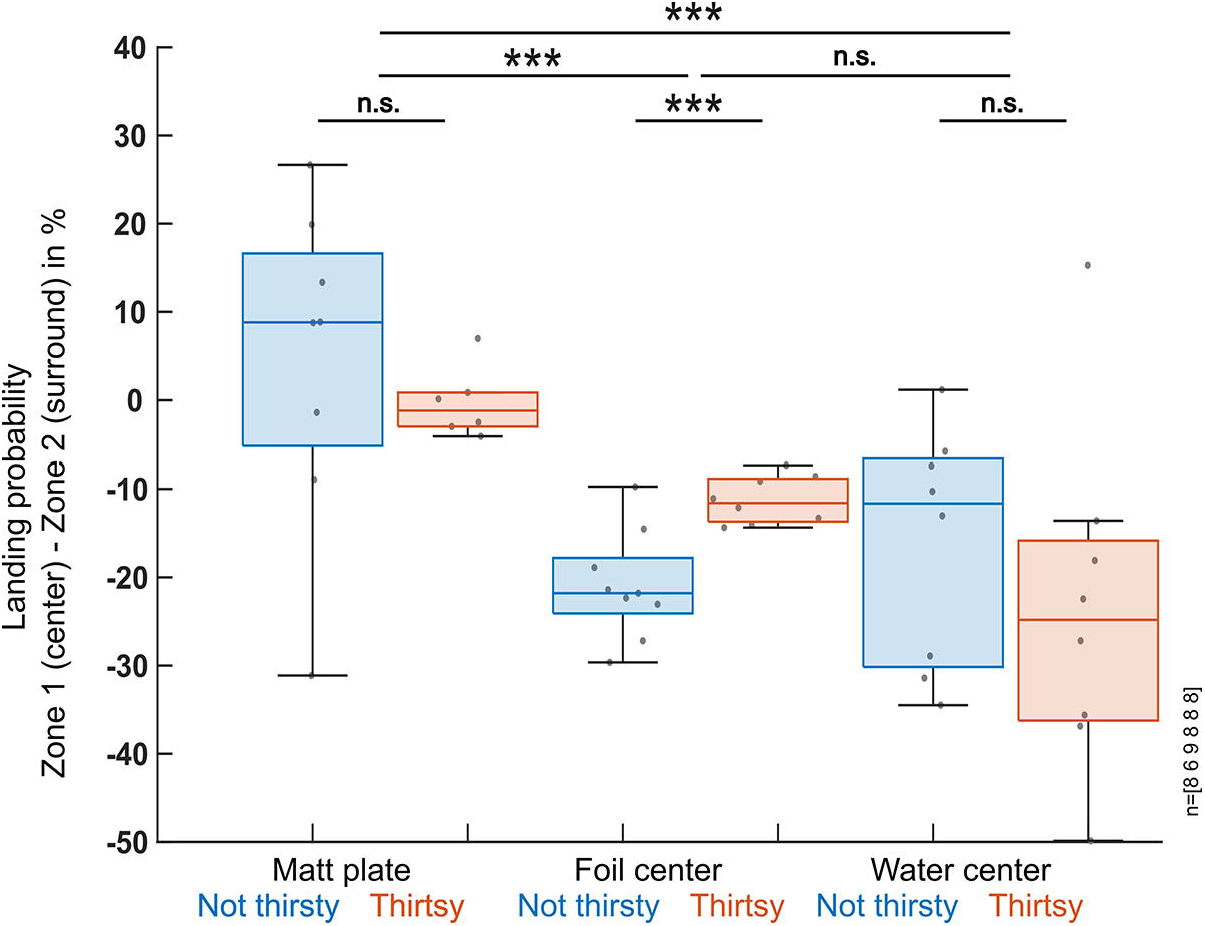
Statistical analysis of landing probabilities across all conditions. Statistical summary of landing probability (zone 1 – zone 2), directly comparing hydrated and dehydrated flies in all three experimental conditions. Number of 60-minute recordings from left to right indicated at the right. *p<0.05, **p<0.01, ***p<0.001.

### Response to water in the darkness

In order to test whether flying flies also interact with the centrally placed water body in the complete absence of any visual cues, we also quantified the spatial distribution of flying flies in darkness with the water stimulus in the center. Indeed, we found that flies showed a slightly higher detection probability in zone 1 as compared to zone 2 (Fig. 7). More importantly, visual inspection of their landing behavior revealed that even in complete darkness, flies would seek out the edge of the water body, where they would then remain immobile for long periods of time (supplemental Fig S1c).

**Figure 7:**
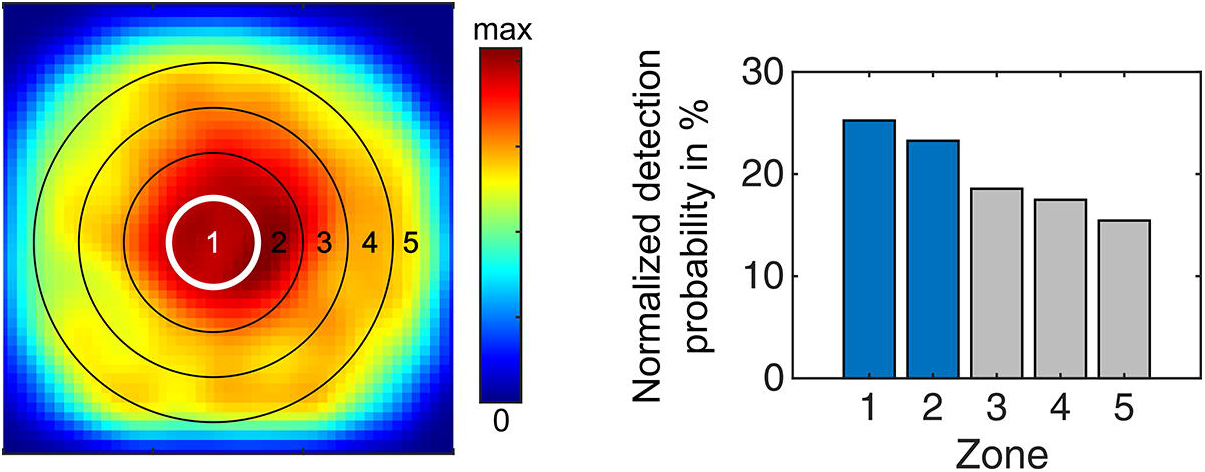
Flight densities of hydrated flies in response to water in complete darkness. Summary of flight densities of hydrated flies in the arena after 60 minutes of recording over a matt plate with a water center in complete darkness. A heatmap of false-colored flight densities is shown, as well as the normalized detection probability for flying *Drosophila* to be in each concentric zone.

## Discussion

The experiments described here used the quantitative analysis of both flight densities and landings and revealed that flies manifest a statistically significant tendency to avoid flying over and landing on shiny surfaces, which are significantly more polarized than the matt surroundings. Significant differences between hydrated and dehydrated flies were only found in the landing probability with a linearly polarizing center (shiny foil). When using water as a reflectant, statistical analysis did not reveal any similar effect for flight densities, neither for thirsty nor hydrated flies. Interestingly, most flies were found in close vicinity to the water body in complete darkness.

Sensitivity to polarized light emanating from the ventral field of view has been described for many other insect species (reviewed by Heinloth et al., 2018 and (Yadav and Shein-Idelson 2021). Although the experimental setup used here is very different from the ones before, the results obtained here are in quite good agreement with these studies, as well as with those using other model systems. While most species show a strong attraction to water/shiny surfaces, only relatively few manifest an avoidance of such stimuli (Shashar, Sabbah et al. 2005).

### A new experimental setup for studying non-celestial polarization vision

The experimental setup used here represents a new, easy-to-use approach for insect behavioral science, enabling the examination of the ventral perception of linearly polarized light using free-flying flies and a robust algorithm for tracking flight maneuvers. Using a population of free-flying *Drosophila* allows for approximating more naturalistic behaviors, yet the setup is much easier to use than comparable 3D tracking approaches (Straw, Branson et al. 2011, Stowers, Hofbauer et al. 2017). Here, a matt acrylic surface was chosen as a negative control, since these materials are known to produce only weakly linearly polarized reflections (Kriska, Bernath et al. 2007, Horvath, Mora et al. 2011). In contrast, shiny transparent overhead foil can induce behavioral responses similar to water seeking (Kriska, Horvath et al. 1998, Wildermuth 1998). In this new assay, large numbers of free flying insects can now be confronted with one crucial (visual) aspect reflected by water (Kriska, Horvath et al. 1998), without providing any non-visual cues usually associated with water.

### Differences in detecting polarized reflections versus water

Characterization of the water stimulus revealed very low values for the degree of polarization, virtually identical to the matt negative control. This was expected, since water had to be placed on a bright surface to insure good contrast for filming and ultimately tracking the flies. However, the white color of such a recipient was previously shown to cause low polarization contrast (Meglic, Ilic et al. 2019). According to these measurements, the DOLP was lowered due to the unpolarized light randomly reflected (scattered) from the white recipient. Hence, the water stimulus most likely served as yet another negative control, by lacking the most important visual aspects of water (shine and polarization), while providing all non-visual qualities of a water source, first and foremost olfactory and gustatory cues (Lin, Owald et al. 2014).

Given that our behavioral paradigm allowed for the presentation of those visual cues associated with water, while omitting the olfactory and gustatory components, it seems plausible to assume that polarized reflections alone induce an overall avoidance response. Flies avoided flying over, and landing on such surfaces. However, in the absence of all visual cues, the olfactory / gustatory components also appear to be sufficient for inducing an avoidance to crash into the water body. This is in good agreement with previous studies demonstrating that flies are able to perceive water non-visually, i.e. using the gustatory sense for measuring the humidity gradient (Ji and Zhu 2015).

### The effect of desiccating flies

After six hours of desiccation, *Drosophila melanogaster*, which under normal conditions are repelled by water, become attracted to it, and this period is sufficient to induce a water-seeking behavior in *Drosophila* (Lin et al., 2014). We only found a difference in landing probabilities between non-thirsty and thirsty flies when confronted with a shiny surface (foil). However, we found no desiccation-based differences in flight distributions. This once again points towards shiny surfaces probably recapitulating only the visual feature of real water, while missing crucial, non-visual features that are necessary for identifying the appetitive or aversive nature of this stimulus. The visual features alone appear to induce the same avoidance response, irrespective of the internal state of the animal.

### The retinal substrate of ventral polarization vision

To the present day, the exact nature of the retinal substrate responsible for *Drosophila* behavioral responses to ventrally presented polarized light remains highly unclear, since no ommatidia reminiscent of the DRA type exist in the ventral part of the adult fly eye (Wada 1974, Labhart and Meyer 1999). Amongst flies, only in horse flies a clear correlation between one of the pale / yellow-like ommatidial types and ventral polarization sensitivity has been found (Meglič et al., 2019), with the untwisted R7H / R8H photoreceptor pairs (harboring horizontally and vertically aligned microvilli in R7 and R8, respectively) being perfectly suited to for polarotaxis (Stavenga et al., 2003; Meglič et al., 2019). Interestingly, in *Drosophila* the same pale inner photoreceptor (R7) subtype is required to induce ventral polarotaxis, but not its yellow counterpart (Wernet et al, 2012). Hence, it remains a possibility that the underlying mechanisms are conserved amongst flies. In contrast, a very different organization of non-DRA ommatidia with polarization-sensitive photoreceptor subtypes was described in another fly species, the Dolichopodid fly *Sympycnus lineatus* (Trujillo-Cenóz and Bernard, 1972).

Future studies should focus on revealing how polarized reflections are detected in the *Drosophila* eye. This knowledge is crucial for understanding how these signals are further processed by the brain, in order to inform behavioral responses. So far, the underlying pathways are understood only for the detection of celestial polarization, in some species (Weir, Henze et al. 2016, Heinze 2017, Omoto, Keles et al. 2017, Timaeus, Geid et al. 2020, Hardcastle, Omoto et al. 2021, Kind, Longden et al. 2021, Sancer and Wernet 2021, Homberg, Hensgen et al. 2022). The behavioral setup we present here now provides an efficient platform for the systematic dissection of the retinal substrate, as well as the underlying circuit elements in *Drosophila melanogaster* (Simpson 2009, Wernet, Huberman et al. 2014).

## Acknowledgements

The authors would like to thank Eugenia Chiappe for providing fly strains, as well as the groups of Robin Hiesinger, Gerit Linneweber, Jochen Pflüger, and Randolf Menzel for valuable input over the years. This work was supported by the AFOSR grant FA9550-19-1-7005 (MFW and GB), by the Deutsche Forschungsgemeinschaft through grants WE 5761/4-1 (MFW) and SPP2205 (MFW), with support from the Fachbereich Biologie, Chemie & Pharmazie of the Freie Universität Berlin (MFW). This work is dedicated to the memory of our friend and colleague Jochen Pflüger.

## Author contributions

MFW and TFM conceived the study. TFM, EJB, AG, ES, and NG collected data. TFM, EJB, AG, ES, NO, NG, GB analyzed the data. MFW, TFM, EJB, and GB wrote the manuscript with input from everyone.

