## Supplemental Table and Figure for "Behavioral responses of free flying *Drosophila melanogaster* to shiny, reflecting surfaces"

**Supplemental Table T1**

| Flight | | | | Landings | | | |
|---|---|---|---|---|---|---|---|
| Not thirsty | | Thirsty | | Not thirsty | | Thirsty | |
| **Number of flies** | **Number of detections** | **Number of flies** | **Number of detections** | **Number of flies** | **Number of landings** | **Number of flies** | **Number of landings** |
| matte | | matte | | matt | | matt | |
| 211 | 280730 | 122 | 722480 | 211 | 95 | 122 | 1000 |
| 150 | 257180 | 119 | 708580 | 150 | 57 | 108 | 1000 |
| 131 | 229850 | 108 | 929900 | 131 | 71 | 47 | 1000 |
| 98 | 204440 | 111 | 923090 | 98 | 101 | 99 | 1000 |
| 149 | 342320 | 106 | 474520 | 149 | 231 | 86 | 1000 |
| 116 | 146270 | 121 | 620160 | 116 | 73 | 110 | 1000 |
| 122 | 323610 | 47 | 238540 | 122 | 243 | | |
| 207 | 467940 | 99 | 674580 | 207 | 160 | | |
| 129 | 408880 | 86 | 772840 | | | | |
| 107 | 603140 | 110 | 671660 | | | | |
| shiny | | shiny | | shiny | | shiny | |
| 111 | 211390 | 127 | 579930 | 111 | 39 | 127 | 1000 |
| 101 | 310170 | 122 | 563460 | 101 | 189 | 122 | 1001 |
| 78 | 181790 | 161 | 717170 | 78 | 230 | 161 | 1000 |
| 85 | 335120 | 179 | 902230 | 85 | 345 | 95 | 1000 |
| 175 | 501990 | 120 | 508490 | 175 | 175 | 115 | 1000 |
| 130 | 311580 | 95 | 456500 | 130 | 127 | 77 | 1000 |
| 105 | 246690 | 115 | 536580 | 116 | 85 | 110 | 1000 |
| 116 | 386540 | 77 | 425320 | 111 | 74 | 113 | 1001 |
| 111 | 335880 | 110 | 553080 | 114 | 250 | | |
| 114 | 510610 | 113 | 485630 | | | | |
| water | | water | | water | | water | |
| 150 | 164810 | 60 | 80077 | 150 | 105 | 60 | 96 |
| 90 | 109890 | 120 | 186660 | 90 | 70 | 120 | 161 |
| 170 | 120410 | 90 | 175020 | 170 | 94 | 90 | 240 |
| 90 | 122970 | 20 | 18588 | 90 | 62 | 20 | 25 |
| 80 | 204750 | 60 | 48488 | 80 | 56 | 60 | 106 |
| 170 | 248450 | 20 | 60042 | 170 | 122 | 20 | 20 |
| 110 | 71571 | 110 | 221280 | 110 | 82 | 110 | 224 |
| 120 | 186860 | 110 | 222620 | 120 | 119 | 110 | 389 |

**Supplemental Figure S1**

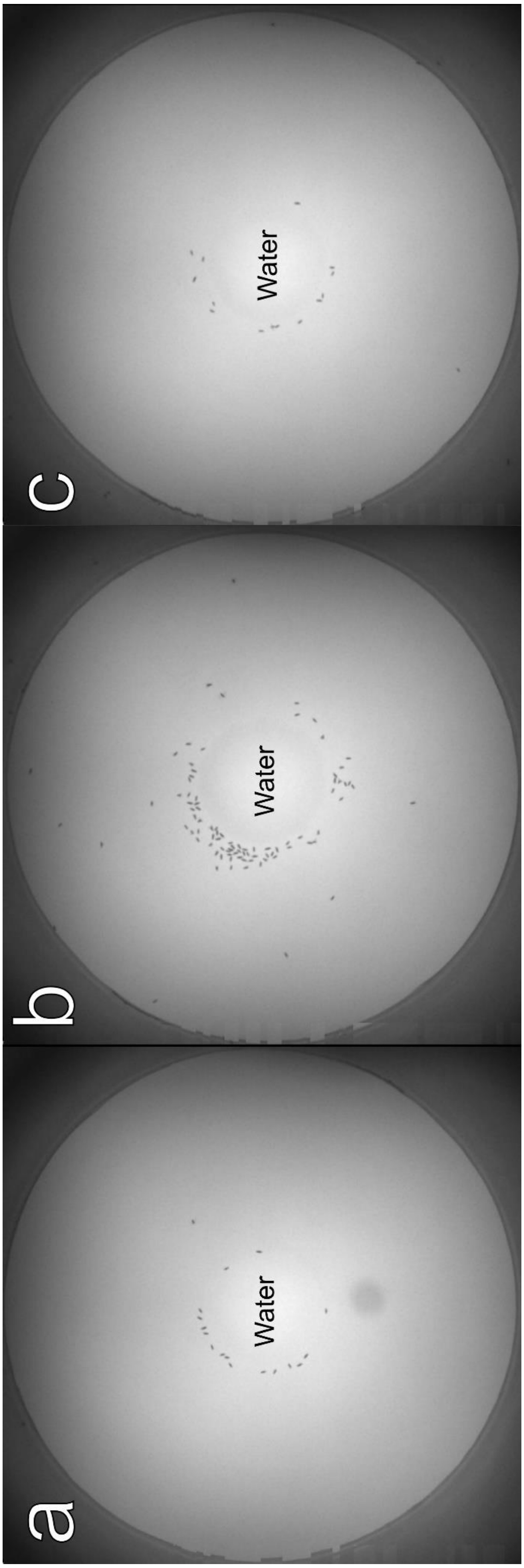

### Supplemental Figure Legends

Supplementary Figure S1: Flies sitting close to the water surface

Different snapshots show the recording of the free-flying wild-type *Drosophila melanogaster* approximately 30 minutes after being released in the arena. The flies were ventrally excited by white light reflected off a water surface. Non-thirsty flies (A), as well as flies dehydrated for six hours (B), and even non-thirsty flies in complete darkness (C), all manifest the same behavior, sitting at the water's edge, suggesting that they use additional sensory cues to detect water sources.
